# The Distribution and Diversity of LuxI/LuxR Quorum Sensing Systems in Proteobacteria (Pseudomonadota)

**DOI:** 10.64898/2026.01.11.698852

**Authors:** Cristina Bez, Marco Sollitto, Vittorio Venturi

## Abstract

In Pseudomonadota (formerly Proteobacteria), the *N*-acyl homoserine lactone (AHL) quorum sensing (QS) system involves LuxI/LuxR modules, where LuxI synthesizes AHLs and LuxR-AHL regulates target gene expression. Despite extensive characterization in many bacterial species, the distribution, genomic organization, and taxonomic comprehensiveness of LuxI/R systems remains unexplored at scale. In this study, we present the first large-scale, genome-wide assessment of LuxI/R QS systems across over 30,000 publicly available genomes at chromosome-level and manually curated spanning 938 genera. Using pfam-based domain annotation, we mapped the distribution, occurrence, and copy number of LuxI/R homologs. LuxI/R systems were identified in ∼32% of surveyed genera, with notable enrichment in symbiotic and plant-associated taxa such as *Rhizobium*, *Burkholderia*, and *Pseudomonas*, and remarkable conservation in pathogenic taxa such as *Yersinia*, *Aeromonas*, and *Acinetobacter*. Conversely, entire genera, including *Escherichia, Salmonella*, and *Klebsiella*, among others, lacked LuxI/R systems across all sequenced strains, suggesting evolutionary loss, niche-specific signaling strategies, or reliance on alternative currently unknown communication systems. Our results also revealed genera with multiple, non-redundant and taxonomically not-related LuxI/R pairs per genome, indicating modular architectures and unpredictable events of horizontal gene transfer events and genetic arrangements. This study delineates the complex distribution and conservation patterns of LuxI/R circuits, providing a resource for future studies into AHL-mediated QS regulation, and microbial community interactions across diverse environments.

**Highlights:**

- LuxI/LuxR quorum-sensing systems show an uneven distribution across Pseudomonadota (formerly Proteobacteria) and evolved through expansion, conservation, and loss, reflecting diverse ecological strategies
- Among ∼31,815 high-quality Proteobacterial genomes, 6,400 (20.1%), corresponding to
- ∼303 genera (∼32%), encoded at least one complete LuxI/LuxR pair.
- Multiple, highly variable LuxI/R copies, up to seven per genome, reveal extensive horizontal gene transfer and mosaic evolutionary histories, especially in rhizobial and plant-associated taxa.
- Phylogenetic analyses distinguish genera with conserved QS architectures from those with fragmented, lineage-independent LuxI/R repertoires, possibly highlighting diverse ecological pressures shaping QS.

## Introduction

Bacteria most often communicate and coordinate community behaviors by releasing signaling compounds through a process known as quorum sensing (QS). This is a contact independent mechanism whereby bacterially produced extracellular signals accumulate in the environment and when they reach critical or “quorum” levels, the expression of target genes is triggered ^1^. Canonical cell-cell QS systems consist of a signaling and sensing module, where the first produces a compound signal, and the second interacts with it and modulates gene expression. Many bacterial species undergo cell-cell communication, each with its own variety of chemical signals that regulate several adaptive phenotypes which increase competitiveness ^2^. The QS response plays a significant role in bacterial pathogenicity and symbiosis as well as facilitating intraspecies, interspecies and interkingdom interactions.

While a variety of QS systems exist, *N*-acyl homoserine lactone-based QS is the best studied and is exclusively found most common system of Gram-negative Pseudomonadota (formerly Proteobacteria). It consists of a LuxI-family *N*-acyl homoserine lactone (AHL) synthase as the signaling module, and a LuxR-family AHL-dependent transcriptional regulator as the sensing module ^3^. Notably, the genes encoding these two modules typically occur as a cognate pair positioned in close genomic proximity, forming an “auto-inducing” regulatory circuit. The first characterized QS system, from which the name LuxI/R originates, was discovered in the marine bacterium *Aliivibrio fischeri*. When *A. fischeri* reaches high cell densities, and consequently, quorum AHL concentrations, within the light organs of certain marine squids, the LuxI/R system regulates bioluminescence, enabling symbiotic light production ^4,5^. Since this discovery, LuxI/R AHL-QS systems have now been experimentally identified and studied in over 70 different bacterial species, including well-studied models such as *Pseudomonas aeruginosa* regulating biofilm and virulence, *Agrobacterium tumefaciens* regulating conjugative transfer of Ti plasmids, *Sinorhizobium meliloti* regulating symbiosis related genes, *Burkholderia* sp. regulating virulence, biofilm and antibiotic production and *Pectobacterium carotovora* regulating plant-degrading enzymes responsible for soft rot disease ^6–8^. Although laboratory studies and simple model systems have revealed much about LuxI/R signaling and regulation, their evolutionary history across Proteobacteria as well as functions in natural and complex microbial communities remains poorly understood.

Genome sequencing studies have revealed that many Pseudomonadota possess canonical LuxI/R QS systems. Surprisingly few minor comparative genomic studies have focused on these systems, for example (i) a 2013 study on *Burkholderia* species demonstrated that LuxI/R systems cluster based on gene topology rather than species taxonomy ^9^, (ii) LuxI/R systems were found to be abundant among proteobacterial members of the *Populus deltoides* root microbiota ^10^, (iii) a comparative study of 62 genomes within the Sphingomonadaceae showed that LuxI/R modules are a common genomic feature in this family ^11^, and (iv) a metagenomic analysis of a nitrifying wastewater community confirmed the presence and distribution of LuxI/R systems in complex environmental assemblages ^12^. Additionally, evolutionary studies based on small genome datasets suggest that LuxI/R regulators originated early in proteobacterial evolution and, in many cases, were acquired through horizontal gene transfer ^13–15^. Despite these specific reports, a comprehensive overview of the distribution, diversity, and evolutionary patterns of LuxI/R systems across all Pseudomonadota is surprisingly lacking. Here, we address this gap through a large-scale *in silico* analysis of more than 30,000 publicly available Pseudomonadota genomes. To our knowledge, this represents the first global and systematic survey of LuxI/R QS systems, revealing their prevalence, diversity, genomic organization, and potential ecological and evolutionary significance across the Pseudomonadota phylum.

## Materials and Methods

### Genomes retrieval and annotation

We retrieved all publicly available complete and chromosome-level Pseudomonadota genomes using the NCBI database (accessed on 12/2024). Genomes were downloaded in FASTA format. Metadata (assembly level, taxonomy, accession) were collected through the NCBI Datasets command-line tool to ensure standardized accession handling. All genomes were annotated using Prokka v1.14.6 ^16^ with default bacterial parameters. Prokka was run to generate standardized GFF3 annotation.

### Identification of LuxI and LuxR homologs

Putative LuxI and LuxR proteins were searched among all predicted proteomes and identified based on PFAM domain signatures using HMMER v3.3 ^17^. Hits were retained if they contained i) PF00765 (Acyl-homoserine-lactone synthase; LuxI) and ii) PF03472 (autoinducer-binding domain; characteristic of LuxR N-terminal receptor domain).

### Removal of false-positive proteins

All candidate LuxI/LuxR sequences were further annotated using InterProScan v5.63 (Jones et al., 2014), which integrates Pfam, InterPro and Gene Ontology domains. Only sequences for which InterProScan independently confirmed the appropriate domain architecture were kept. Domain boundaries, catalytic residues, and HTH-DNA-binding motifs were manually inspected when necessary. To validate homology, each candidate LuxI/LuxR sequence was searched against the NCBI non-redundant (nr) protein database using DIAMOND BLASTp ^18^. Searches were performed and kept with an E-value threshold of 1e-5, minimum identity of 30%, and coverage ≥70%. The top-scoring hits were used to confirm subfamily identity and exclude false positives.

### Complete-system identification and phylogenetic analysis

To determine whether each genome encoded a complete quorum-sensing system, we used the genomic position of LuxI as an anchor. For every identified LuxI homolog, we retrieved its genomic coordinates from the Prokka GFF3 annotation file and searched for the nearest LuxR homolog. The intergenic distance between the two loci was calculated as the absolute difference between the coordinates of the LuxR and LuxI coding sequences. Pairs with a LuxR-LuxI distance ≤15 kb were classified as complete LuxI/LuxR systems. This distance threshold follows the typical genomic organization of co-occurring LuxI/LuxR operons. The resulting dataset represents high-confidence LuxI/LuxR homologs suitable for downstream phylogenetic and comparative analyses. Finally, the concatenated LuxI-LuxR complete system was initially aligned with ClustalO ^19^, poorly aligned regions were removed using trimAl ^20^. The resulting alignment was then used to infer the phylogenetic tree with IQ-TREE ^21^. The phylogenetic tree was visualized using EMPress^22^.

## Results and Discussion

### LuxI/LuxR quorum sensing systems are unevenly distributed and conserved across Proteobacterial lineages

We used Pfam domains defining the *N*-acyl homoserine lactone (AHL) producing and responding LuxI and LuxR QS protein cognate pair in order to search a comprehensive, well-curated, and high-quality set of Proteobacterial (now reclassified as Pseudomonadota) genomes, generated from publicly available data in the NCBI database. Although the phylum has been recently renamed and the former Beta- and Gammaproteobacteria have been merged into a single Gammaproteobacteria class, the genomes were taxonomically annotated in NCBI under the previous classification (Alpha-, Beta-, and Gammaproteobacteria). Therefore, we refer to these original designations throughout the manuscript. Our dataset comprised ∼31,815 genomes, representing 938 genera. Among these, 6,400 genomes (20.1%), around one-third of all the genera (∼303; ∼32%), encoded at least one complete LuxI/LuxR pair. The relative frequency of LuxI/LuxR-positive genomes for classes was 41.7% in Alphaproteobacteria, 24.8% in Betaproteobacteria, and 16.6% in Gammaproteobacteria (Figure 1). These proportions suggest that LuxI/R systems are more conserved and functionally relevant within Alphaproteobacteria, whereas Gammaproteobacteria display greater regulatory diversification and evidence of partial system loss.

**Figure 1:**
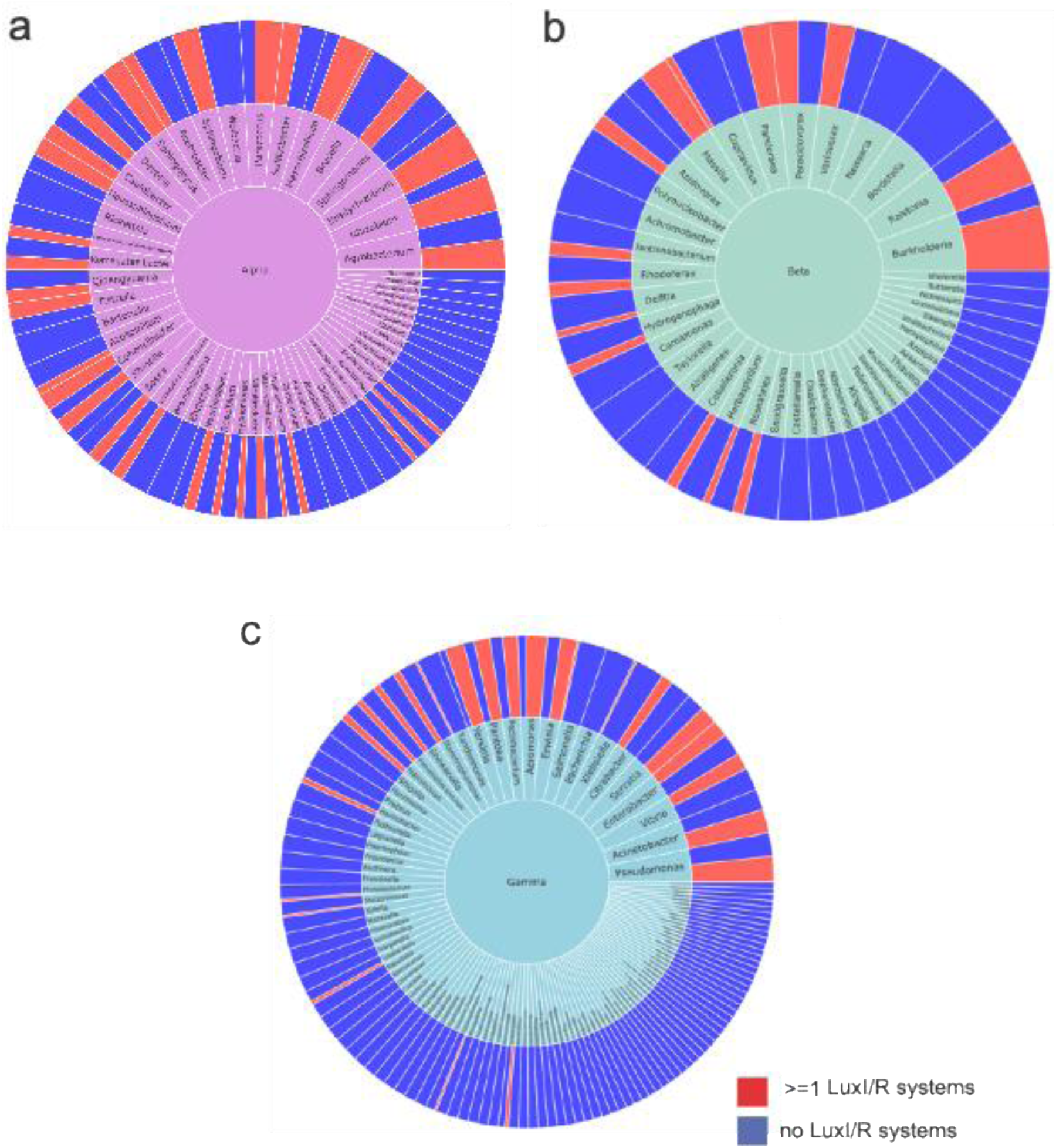
Distribution and number of LuxI/R QS systems among a) Alphaproteobacteria, b) Betaproteobacteria and c) Gammaproteobacteria genomes. The inner rings represent taxonomic hierarchy at Genus level, while the outer ring shows genome counts (log10) with blue color indicating genomes lacking QS systems, and red genomes carrying one or more complete LuxI/LuxR systems. Genera with at least 50 genomes in the dataset were included in this figure. A complete overview of all analyzed genera is available at the following link: https://github.com/marco91sol/LuxI-LuXR_Pseudomonadota/tree/main/sunburst_by_class.

Within the Alphaproteobacteria, seven genera were predominant, collectively harboring the highest number of LuxI/LuxR systems (Figure 1a). The proportion of genomes containing at least one AHL-QS system was exceptionally high across most of these genera, 100% in *Pseudosulfitobacter* and *Phaeobacter*, 98.9% in *Sinorhizobium*, 96.9% in *Bradyrhizobium*, 93.3% in *Rhizobium*, and 92.8% in *Mesorhizobium*, indicating that QS regulation is an almost universal feature within these lineages. Only *Agrobacterium* showed a lower frequency (54.3%), displaying variability among the analyzed species and genomes.

Among the Betaproteobacteria, the distribution of LuxI/LuxR systems was dominated by four genera, comprising approximately 84% of all betaproteobacterial genomes (Figure 1b). The proportion of genomes encoding a QS system was consistently high within these taxa, 98.6% in *Burkholderia*, 96.6% in *Paraburkholderia*, 96% in *Chromobacterium*, and 84.8% in *Ralstonia*, suggesting a strong evolutionary conservation and functional relevance of AHL-QS within these genera.

When focusing only on the genera with the largest number of genomes available in the dataset, the proportion of LuxI/LuxR-positive genomes among gammaproteobacteria, normalized to the total genomes per genus, was 97.5% in *Yersinia*, 97.2% in *Aeromonas*, 70.3% in *Acinetobacter*, 62.1% in *Pseudomonas*, and 51.8% in *Serratia* (Figure 1c). This suggests that the presence of at least one LuxI/R system is a common and conserved trait in *Yersinia* and *Aeromonas*, whereas the absence of such systems (both approximately 50%) is equally frequent among *Serratia* and *Pseudomonas*. A closer analysis at the species level revealed that *Yersinia rochesterensis* was the only species lacking any complete LuxI/LuxR systems among *Yersinia* genomes. In contrast, among *Serratia* genomes, *Serratia marcescens* exhibited high intra-species variability, with approximately half of the analyzed genomes containing one AHL-QS system and the remaining half lacking it entirely. Among *Pseudomonas* species, *P. aeruginosa* consistently harbored two systems, whereas *P. fluorescens* displayed a heterogeneous pattern, with the majority of genomes lacking complete LuxI/LuxR pairs.

Interestingly, only four genera showed a complete retention of LuxI/R QS systems, with every analyzed genome belonging to different species, encoding at least one system. Among these genera, *Edwardsiella* (166 genomes; 6 species) and *Dickeya* (98 genomes; 12 species), are facultative pathogens of fish/humans and plants, respectively, while *Pseudosulfitobacter* (61 genomes; 2 species ) and *Phaeobacter* (50 genomes; 3 species) are both marine bacteria symbiotically associated with algae and other marine organisms. This finding suggests that in these proteobacterial lineages, AHL-QS has been evolutionarily conserved as a core regulatory mechanism possibly playing a role in host interaction, virulence regulation or another group behaviour. Overall, the taxonomic diversity of QS LuxI/R system distribution highlights the uneven presence of QS systems, pointing to varied ecological roles and regulatory adaptations across Proteobacteria. The full distribution of LuxI/R systems across all genera is provided as an interactive sunburst plot, and it is deposited in the public GitHub repository page https://github.com/marco91sol/LuxI-LuXR_Pseudomonadota/tree/main/sunburst_by_class.

### Expansion and variability of multi-system LuxI/LuxR QS in Proteobacteria

We detected several genomes encoding more than one, and up to seven, archetypal LuxI/LuxR systems across all three bacterial classes, although multi-system occurrences were overwhelmingly concentrated in Alphaproteobacteria (Figure 2a). Within this class, high-copy numbers were primarily found among rhizobial taxa (e.g., *Bradyrhizobium*, *Mesorhizobium*, *Rhizobium*) and related genera such as *Paracoccus*, *Methylobacterium*, and *Sphingobium*. The presence of up to seven systems in some *Rhizobium* strains evidences the exceptional evolutionary plasticity of QS in this group and its likely central role in regulating symbiosis and group-associated processes. Figure 2a summarizes the top genera categorized by the number of LuxI/R systems (one, two, three, or more than four) encoded in their genomes. Although the number of available genomes per genus influenced these counts, the dataset offers a representative overview of LuxI/R system diversity. Genera predominantly carrying a single LuxI/R system (*Acinetobacter, Aeromonas, Dickeya, Serratia, Enterobacter, and Chromobacterium*) likely rely on well conserved AHL-based QS circuits for some coordinated group gene regulation (see below). Those with two systems (*Pseudomonas, Yersinia, Erwinia, and Pseudosulfitobacter*) may utilize multiple QS pathways for a more refined regulatory control, whereas genera with three or more systems (*Burkholderia, Pseudomonas, Rhizobium, Bradyrhizobium, Mesorhizobium*) possibly possess a more complex hierarchy and AHL-QS architectures, larger regulons, host–microbe interactions, high genetic plasticity and environmental survival strategies.

**Figure 2:**
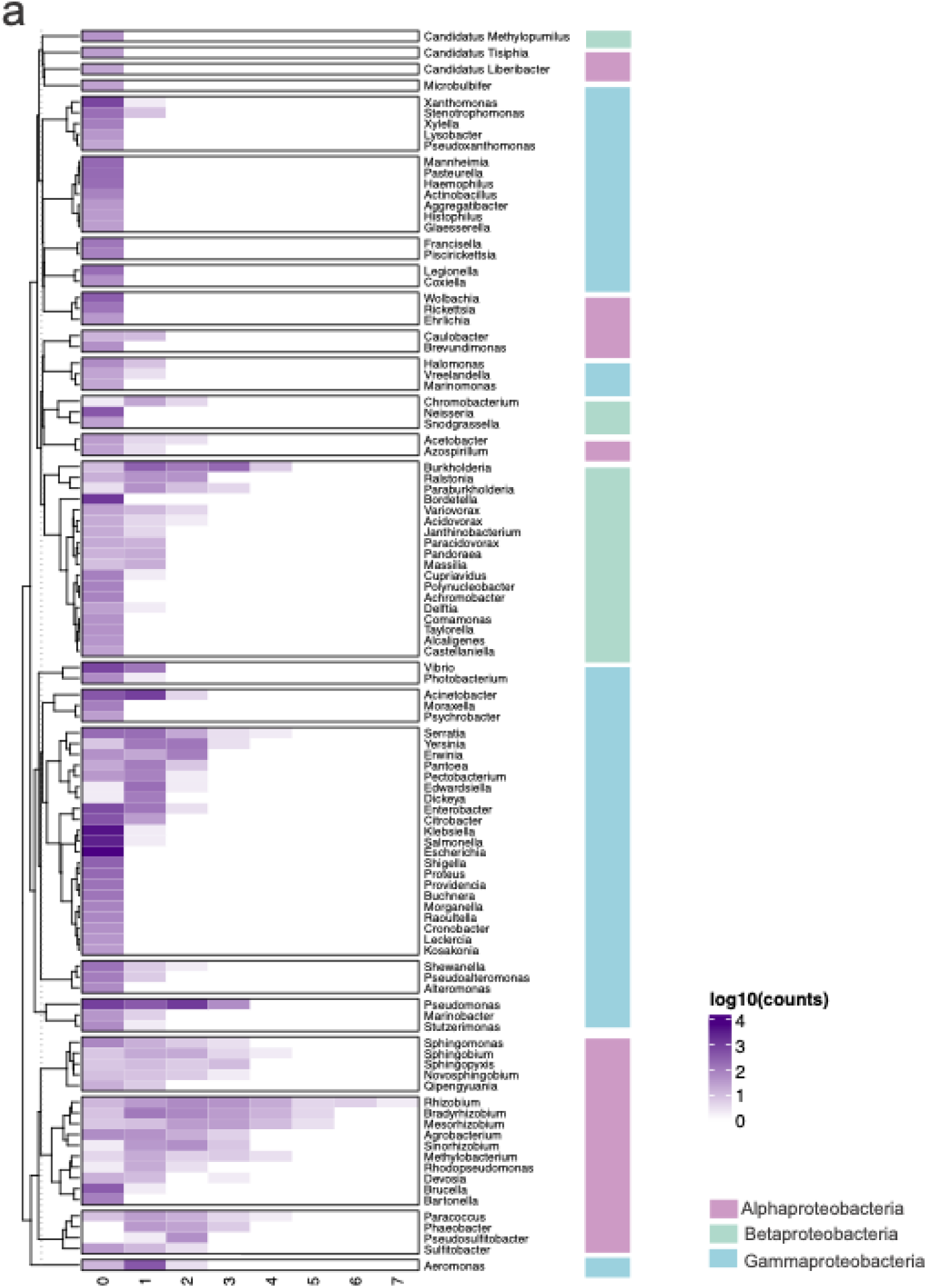
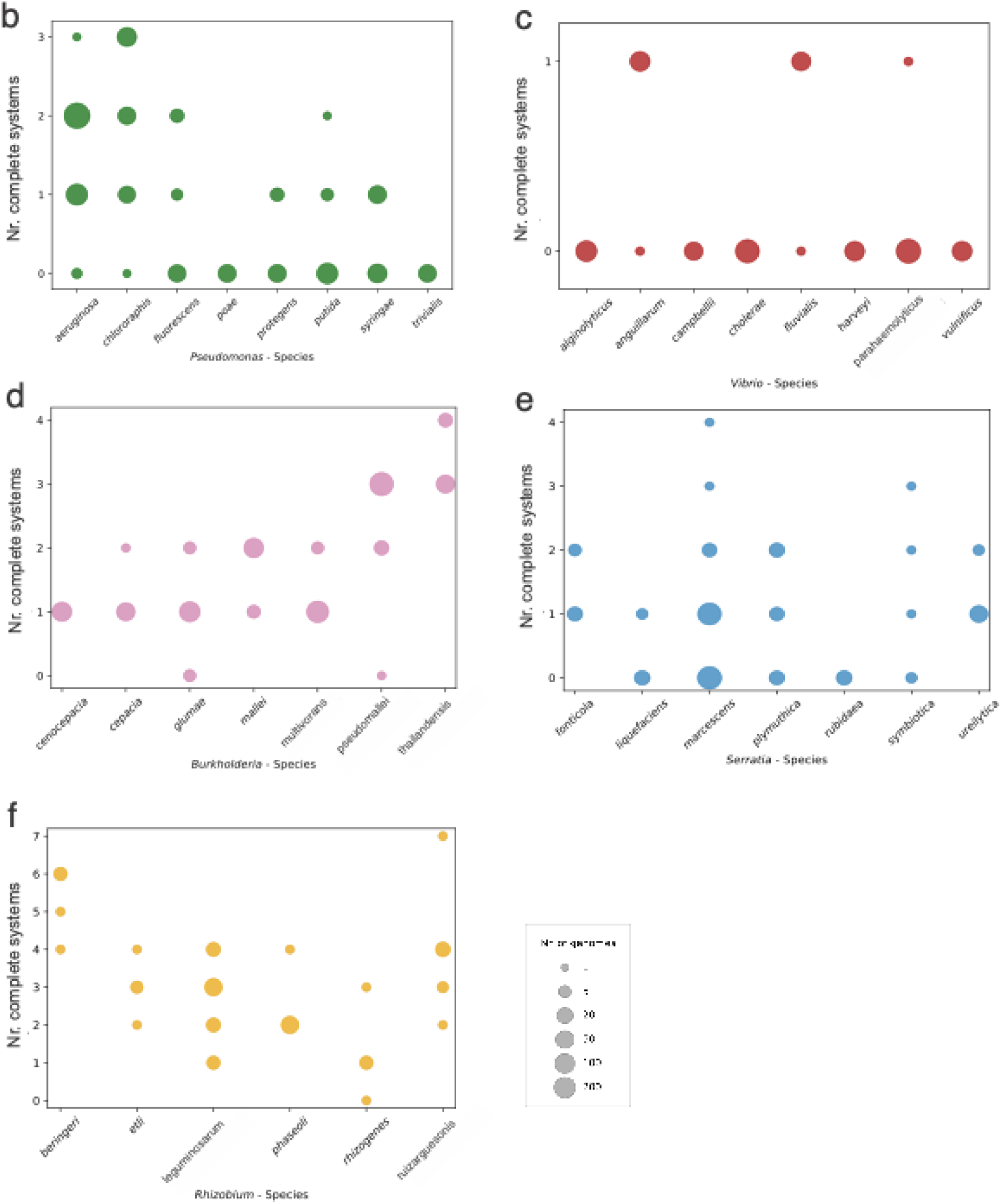
Variability of multi-system LuxI/R in Pseudomonadota. a) Heatmap showing the number of complete QS systems for the most representative genera, arranged and colored by class. Bubble plots showing the number of complete LuxI/LuxR systems identified per species within five representative genera (b) Pseudomonas, (c) *Vibrio*, (d) *Burkholderia*, (e) *Serratia*, and (f) *Rhizobium*. Bubble size is proportional to the number of genomes analyzed for each species, ranging from 1 to over 200 genomes, as indicated in the legend. These plots highlight the marked heterogeneity in LuxI/LuxR system abundance across species within the same genus, reflecting the diverse evolutionary trajectories and ecological strategies of Proteobacteria.

Interestingly, several genera such as *Pseudomonas, Vibrio, Burkholderia, Serratia* and *Rhizobium*, among others, displayed considerable intra-species variability, with some species, or even individual strains, harboring one and up to four LuxI/R systems while others lacked them entirely. This intra-species and inter-species heterogeneity likely reflects a combination of horizontal gene transfer, gene loss, or lineage-specific regulatory adaptation (Figure 2b-f). For instance, within the genus *Serratia*, *S. marcescens* exhibited substantial intra-species variability, with individual strains encoding from zero to four LuxI/LuxR systems, whereas all analyzed strains of *S. rubidaea* uniformly lacked any AHL-QS systems (Figure 2e). A similar pattern was observed in *Burkholderia*: strains of *B. glumae* encoded between zero and two systems, while *B. thailandensis* consistently carried between three and four (Figure 2d). Notably, *Rhizobium* also showed pronounced species-level heterogeneity. All examined strains of *R. beringeri* harbored four to six complete systems, in contrast genomes of *R. leguminosarum* strains contained between one and four systems (Figure 2f).

Together, these examples highlight the mosaic distribution of AHL-QS systems across Proteobacteria, being likely shaped by species-specific ecological pressures, genome plasticity, and symbiotic or pathogenic lifestyles.

### A large fraction of Proteobacterial genera lack or have lost canonical LuxI/LuxR systems

Our findings indicated that the vast majority (around two-thirds) of the Proteobacterial genera, and consequently most genomes, do not possess a canonical LuxI/R QS system. In particular, ∼640 genera completely lacked an AHL-QS system, a pattern that appears consistently maintained across all the genomes investigated. These genera occupy diverse ecological niches and display a wide range of host-environment interactions. They include opportunistic pathogens of humans and animals such as *Escherichia*, *Klebsiella*, *Salmonella*, *Shigella*, *Proteus*, *Providencia*, *Legionella*, *Moraxella*, *Haemophilus*, *Pasteurella*; phytopathogens such as *Xanthomonas* and *Xylella*; obligate symbionts and intracellular bacteria like *Buchnera* and *Wolbachia*; pathogenic intracellular parasites of animals and humans, including *Rickettsia*, *Brucella*, *Bartonella*, and *Francisella*; and finally, free-living aquatic bacteria such as *Alteromonas*, *Polynucleobacter*, and *Cupriavidus*, which inhabit marine or freshwater environments. This absence could be explained by several, non-mutually exclusive factors: (i) ecological specialization, as observed in insect endosymbionts such as *Buchnera* and *Wolbachia*, which have highly reduced genomes encoding only the genes necessary for symbiosis ^23^; (ii) replacement by alternative signaling mechanisms, as in *Xanthomonas* and *Xylella*, which rely on DSF and DSF-like cell–cell signaling systems ^24^, or in *Legionella*, which employs the Lqs system (LqsA/LqsR/LqsS) based on α-hydroxy acid signals ^25^; and (iii) evolutionary loss of canonical components, as in *Escherichia*, *Salmonella*, and *Klebsiella*, which have lost the *luxI* gene but retained *sdiA*, a LuxR-solo homolog capable of sensing exogenous AHLs ^26^.

Overall, these patterns reveal that the absence of canonical LuxI/R is not an exception but a widespread and recurrent evolutionary strategy across bacteria with highly diverse lifestyles.

### Phylogenetic relationships of LuxI/R systems reveal lineage-specific signatures and mosaic evolution

To investigate how the primary structure of LuxI/R AHL-QS systems relate to each other, we constructed a phylogenetic tree based on concatenated LuxI-LuxR protein sequences. This approach allowed us to evaluate not only the presence and number of systems within each genome, but also their degree of sequence conservation across taxa.

A clear pattern emerged for several genera in which the architecture of the AHL QS network is simple, typically one LuxI/R system which was highly conserved. For example, all examined strains of *Acinetobacter, Aeromonas, and Edwardsiella* harbored a single LuxI/R cognate pair which was highly similar across strains. Such conservation suggests that these systems are genus-level signatures and likely tied to their ecological adaptation and lifestyle (Figure 3). In genera with more complex QS architectures, the phylogeny also reflects well-defined organization. *P. aeruginosa* is a clear example: it typically encoded two systems called LasI/R and RhlI/R ^27^. In the phylogenetic tree, all Las-type systems clustered together in a tight, conserved clade, while all Rhl-type systems formed a separate and equally conserved clade (Figure 3, green color of the external ring). This pattern indicates that they belong to the core or pan-genome of *P. aeruginosa*, with distinct evolutionary lines (Figure 3). A similar situation was visible in *Burkholderia*, where the three LuxI/R systems called CepI/R, BtaI/BtaR and BviI/BviR ^28^ formed structured clusters, again consistent with long-term vertical inheritance (Figure 3, violet color of the external ring) and multi-layered signalling.

**Figure 3.**
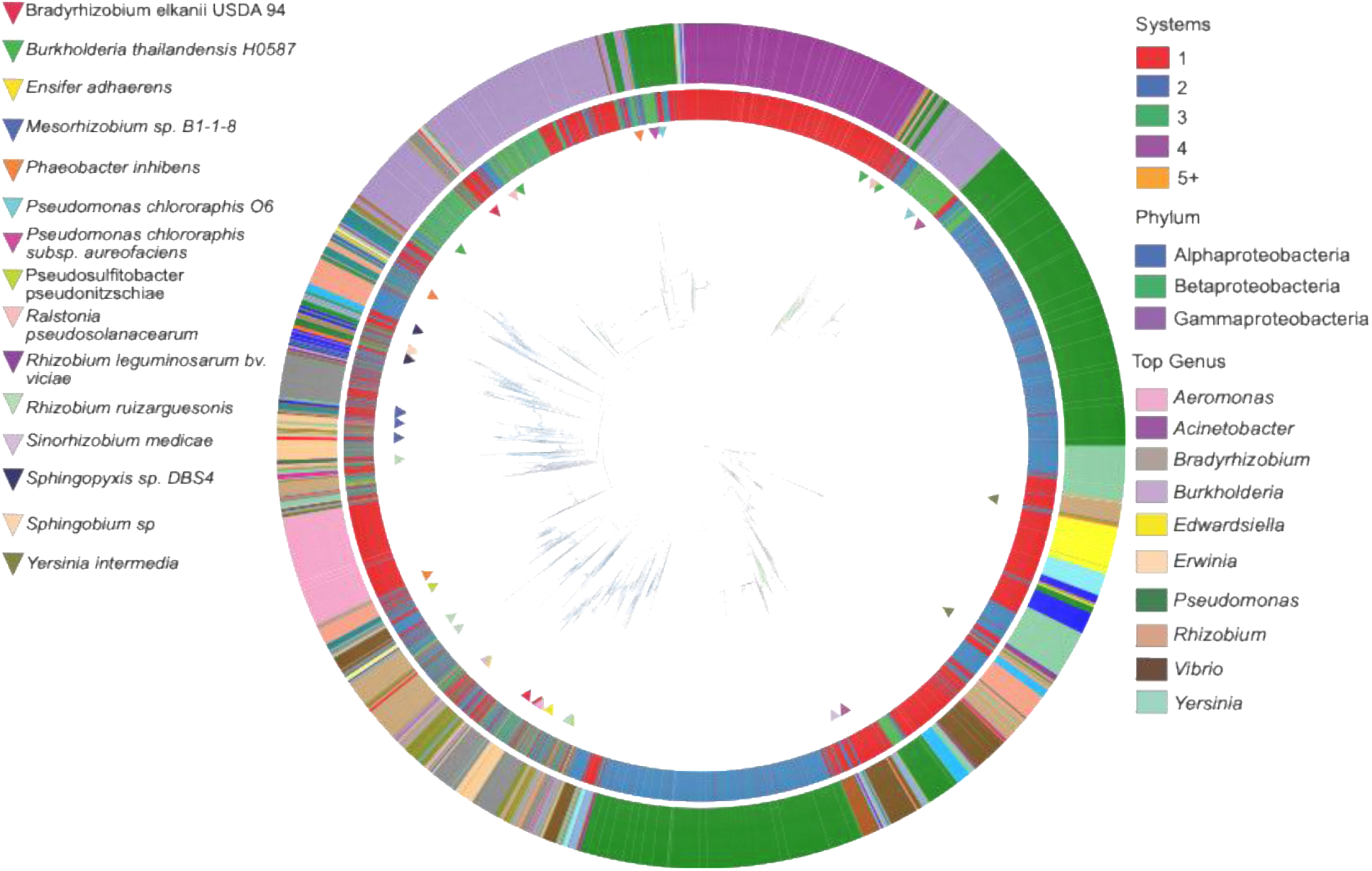
Phylogenetic distribution of LuxI/R QS systems across Pseudomonadota. Maximum-likelihood phylogeny showing the diversity and genomic abundance of LuxI/R systems. Outer rings indicate (i) Genera and in the legend are highlighted the most representative one number, and (ii) number of LuxI/R systems per genome. The branches of the tree are colored based on the phylum-level taxonomy. Example of genomes with exceptionally high number of complete systems are annotated with colored triangles around the tree. The tree is visualized and annotated using EMPress.

In contrast, the rhizobial genera displayed the opposite trend. Species of *Rhizobium* possessed multiple LuxI/R systems with very low sequence similarity, scattered across distant branches of the tree. This scattering pattern is indicative of weak conservation across strains and substantial horizontal gene transfer, producing a mosaic set of QS systems. A striking example was *Bradyrhizobium elkanii USDA 94* (red triangles in Figure 3) and *Sinorhizobium medicae* (violet triangles). These species share one LuxI/R system type, consistent with a common evolutionary origin, yet each also encodes additional systems that fall into entirely different phylogenetic clusters. This reinforces the idea that rhizobial QS systems have been shaped by extensive gene turnover, acquisition, and exchange.

A similar mosaic signature appeared in *Phaeobacter inhibens* (highlighted as an orange triangle in Figure 3), which contained three systems with no apparent homology to one another. One system clustered tightly with the canonical *Aeromonas*-like type, while another group with one of the major *Burkholderia* QS systems. This result suggests that its QS repertoire belongs to the accessory genome and was likely assembled through repeated HGT events. Finally, the situation in *Ralstonia pseudosolanacearum* and *B. thailandensis H0587* is particularly interesting. The two LuxI/R systems in *Ralstonia* showed high homology to two of the four systems in *B. thailandensis*, pointing toward possible niche overlap, ecological interaction, or shared metabolic constraints that favor the maintenance of similar signaling AHL QS systems possibly with similar functional roles.

## Conclusions

LuxI/LuxR QS systems are not uniformly conserved across Proteobacteria. Their prevalence varies both across classes and within genera and even species, indicating multiple lineage-specific evolutionary trajectories, mainly resulting in expansion, conservation and loss or replacement of canonical QS systems. Some genera have undergone significant expansion of QS modules, which correlates with competitive environments that require regulatory plasticity. On the other hand, some genera have lost them entirely, while some retain only the systems that are strictly necessary. Based on our results, the number and types of QS systems likely align with specific ecological strategies: (i) symbiotic and plant-associated bacteria often display expanded and diversified QS repertoires; (ii) pathogens tend to retain conserved, vertically inherited systems; (iii) intracellular or highly specialized taxa frequently show near-complete loss of canonical QS elements; and (iv) marine mutualists and opportunistic pathogens typically maintain stable, strongly conserved QS systems. Beyond system number and simple presence/absence patterns, phylogenetic analyses show that the distribution of QS systems often do not follow species phylogeny. Several taxa display extreme mosaicism, with LuxI/R homologs scattered across distant clades. This supports the view that, in many species, LuxI/LuxR pairs function as accessory regulatory modules, which are advantageous under specific conditions but not universally required. In fact, approximately two-thirds of Proteobacterial genera do not possess LuxI/LuxR systems. This widespread absence indicates that AHL-based QS is not universal and is often evolutionarily dispensable. Losses correlate with specialized lifestyles, such as intracellular symbiosis, obligate parasitism, host-type restriction, or niche-specificity. This compendium of LuxI/LuxR distribution across proteobacterial lineages provides multiple avenues for future investigation, including: (i) empirical tests to determine whether different genera with highly conserved QS systems use their systems to regulate comparable core functions; (ii) in strains carrying multiple QS modules, clarifying whether these systems operate hierarchically, redundantly, or in response to distinct environmental cues; (iii) surveying LuxI/LuxR presence, abundance, and activity in environmental metagenomes to assess whether patterns observed in isolate genomes mirror community-level dynamics and (iv) investigating the types of AHLs produced across genera/species and how these signal chemistries relate to each lineage’s environment and lifestyle.

All of these findings raise important questions about the roles of QS in environmental strains and multispecies communities, contexts that remain underexplored and warrant further investigation.

## Acknowledgements

C.B. benefits funding from the European Union’s Horizon Europe research and innovation programme under the Marie Skłodowska-Curie grant agreement No 101150379.

## Authors contributions

C.B. (Conceptualization, Data curation, Formal Analysis, Investigation, Visualization, Writing – original draft, Writing – review & editing), M.S. (Data curation, Formal Analysis, Methodology, Investigation, Visualization, Writing – original draft, Writing – review & editing), V.V. (Conceptualization, Funding acquisition, Investigation, Project administration, Resources, Supervision, Validation, Writing – original draft, Writing – review & editing).

## Data availability

All data supporting this analysis are publicly available on GitHub page: https://github.com/marco91sol/LuxI-LuXR_Pseudomonadota/tree/main

